# *De novo* design of buttressed loops for sculpting protein functions

**DOI:** 10.1101/2023.08.22.554384

**Authors:** Hanlun Jiang, Kevin M. Jude, Kejia Wu, Jorge Fallas, George Ueda, TJ Brunette, Derrick Hicks, Harley Pyles, Aerin Yang, Lauren Carter, Mila Lamb, Xinting Li, Paul M. Levine, Lance Stewart, K. Christopher Garcia, David Baker

**Affiliations:** Department of Biochemistry, University of Washington; Institute for Protein Design, University of Washington; Howard Hughes Medical Institute, Stanford University School of Medicine; Biological Physics, Structure and Design Graduate Program, University of Washington; Department of Molecular and Cellular Physiology, Stanford University School of Medicine; Department of Structural Biology, Stanford University School of Medicine; Howard Hughes Medical Institute, University of Washington

## Abstract

In natural proteins, structured loops play central roles in molecular recognition, signal transduction and enzyme catalysis. However, because of the intrinsic flexibility and irregularity of loop regions, organizing multiple structured loops at protein functional sites has been very difficult to achieve by *de novo* protein design. Here we describe a solution to this problem that generates structured loops buttressed by extensive hydrogen bonding interactions with two neighboring loops and with secondary structure elements. We use this approach to design tandem repeat proteins with buttressed loops ranging from 9 to 14 residues in length. Experimental characterization shows the designs are folded and monodisperse, highly soluble, and thermally stable. Crystal structures are in close agreement with the computational design models, with the loops structured and buttressed by their neighbors as designed. We demonstrate the functionality afforded by loop buttressing by designing and characterizing binders for extended peptides in which the loops form one side of an extended binding pocket. The ability to design multiple structured loops should contribute quite generally to efforts to design new protein functions.

## Introduction

While antibodies still play central roles in protein therapeutics, progress has been made in drug development using non-antibody binding proteins which show superior properties in thermal/pH stability, binding affinities, tissue delivery and industrial-scale manufacture^1–3^. The two main approaches are random library selection methods and computational protein design. Perhaps the most successful scaffold for random library selection has been the ankyrin repeat^4,5^; the Plückthun group has built libraries of Designed Ankyrin Repeat Proteins (DARPins) which have been used to identify high-affinity binding proteins via high-throughput screening methods which have had multiple successes in preclinical studies^3,5,6^. Ankyrin repeat proteins have a repeating architecture with structured, hairpin-shape loops extending from the helices to an extended binding groove that is geometrically compatible with many globular protein targets. Despite these successes, the global shape diversity of DARPins is limited by the use of a single base scaffold. Computational design of binding proteins does not have this limitation as a very wide range of scaffolds can be used, with shapes more optimal to bind the target protein of interest. However, this advantage thus far has come with a different limitation: because of the inherent flexibility and lack of extensive backbone hydrogen bonding of long loop regions, protein binder design has focused on scaffolds and binding sites primarily composed of alpha helical^7^ or beta strand^8^ secondary structure, which has limited the achievable local shape diversity.

## Results

Here we set out to overcome the challenges in de novo design of long loops on the one hand, and limitations of ankyrin scaffolds in global shape diversity on the other, by computationally designing repeat proteins with multiple long loops buttressed by loop-loop interactions (Fig. 1A). To achieve this goal, we divided the problem into two subproblems: first, the generation of repeating scaffold backbone conformations compatible with loop buttressing, and second, the generation of loop backbone conformations compatible with a dense network of hydrogen bonds and hydrophobic interactions between pairs of loops, and between the loops and the underlying scaffold.

**Fig. 1.**
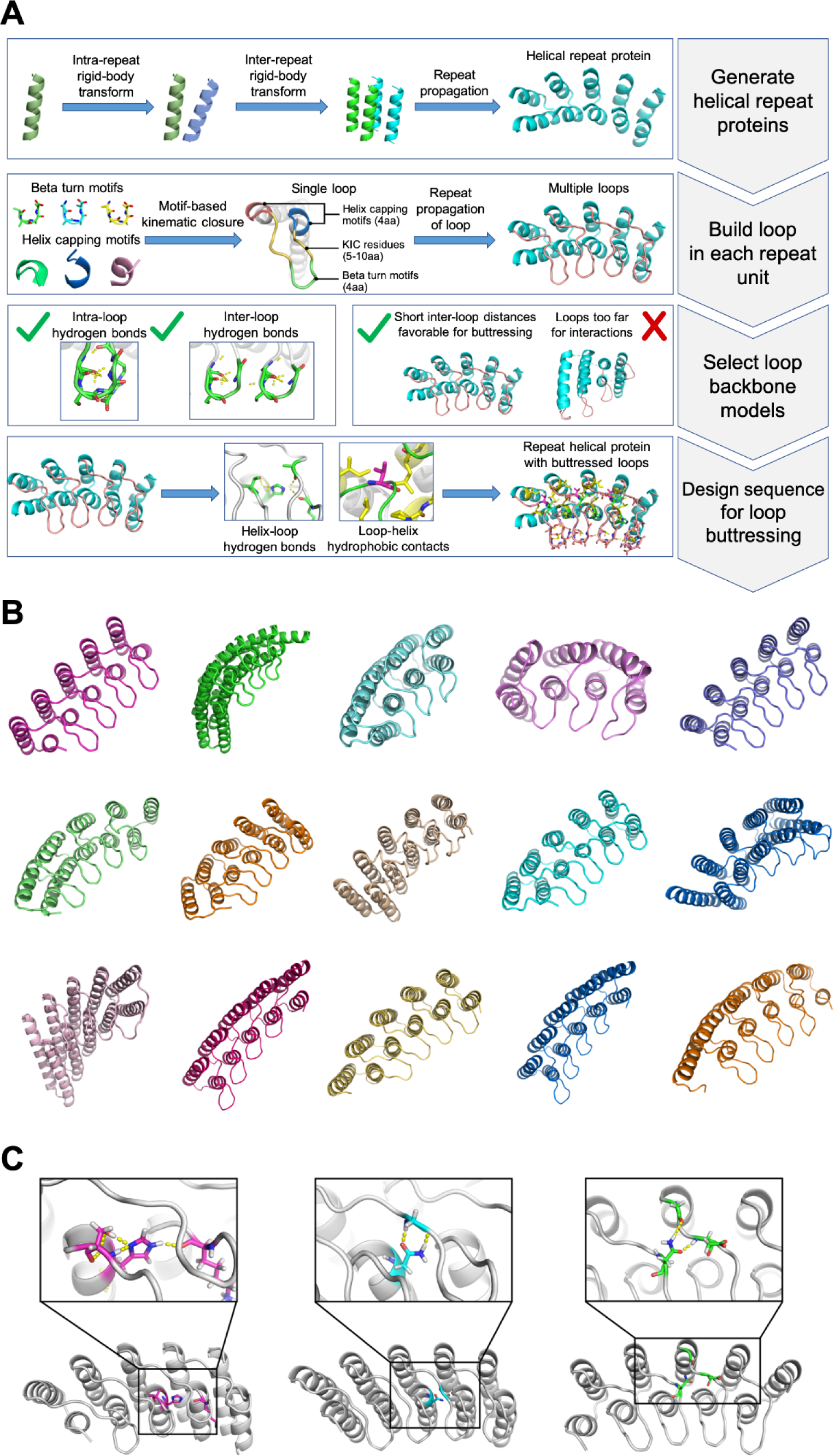
Computational design of repeat proteins with buttressed loops. **(A)** Design strategy for generating and stabilizing multiple loops in helical repeat proteins. **(B)** A gallery of diverse designed proteins that pass the *in silico* design filters. **(C)** Loop buttressing bidentate hydrogen bonds in the designed proteins.

We developed a computational method for generating a wide range of repeat protein backbones geometrically compatible with the insertion of long loops (Fig. 1, top row). Previous approaches of designing helical repeat proteins have utilized fragment assembly methods to assemble repeat units with short loops connecting them^9^. While these methods can generate considerable diversity, we found that they did not provide sufficient control over backbone positions for designing loop buttressing. Instead, we developed a parametric repeat protein generation method which enables precise control over backbone placement. We generated diverse repeat units consisting of two idealized helices by systematically sampling the lengths of the helices (from 12 to 28 residues) and the six rigid-body degrees of freedom between the two helices. We next sampled the six rigid-body degrees of freedom between repeat units, and applied the same transform repeatedly to generate a disconnected repeat protein model. Finally, we connect pairs of sequence-adjacent helices using a native protein based loop lookup protocol that grafts on the 3-6aa loop that best fits onto the termini of the helices^10^. In extensive model building experiments, we found that to enable installation of long loops onto these parametrically generated models, the termini of the helices had to be less than 18Å apart, and we removed backbone models where the distance between termini was greater than this value. We also eliminated poorly packed models with fewer than 28% of the residues in a buried core. The resulting repeat protein models have well-defined core regions and are slightly curved with little or no twisting between the repeat units.

We next sought to develop a general method for building multiple long loops that buttress one another onto protein scaffolds (Fig. 1, second and third rows). Surveying the structured, long loops in natural proteins, we observed that these frequently contain beta turns with strand-like hydrogen bonds flanking the turn residues which contribute to stabilization of the specific loop conformation. We also observed that natural proteins often employ helix capping interactions between sidechain or backbone on the loop residue and the backbone of the helix from which it emanates; these help specify the orientation of the loop as it leaves the helix. Based on these observations, we constructed and curated libraries of beta turn motifs and helix capping motifs by clustering four-residue native protein fragments, and selected the clusters that fulfilled the requirements of hydrogen bonds as described in the Methods. During the loop sampling, these motifs were randomly selected and incorporated in a single loop growing from the N-terminus of a helix. Using generalized kinematic closure^11^, we then connected the C-terminus of the loop to the next helix in the backbone model. The resulting loop was then propagated to each repeat unit to generate a complete repeat protein model with multiple long loops. To specifically favor loops which could be buttressed with hydrogen bond networks, we required that models have at least two intraloop backbone-to-backbone hydrogen bonds within each repeat unit, and at least one interloop backbone-to-backbone hydrogen bond between the repeat unit neighbors. To favor interactions between the long loops and the helices, we further filtered the models by requiring at least five residues within 8Å of the closest helical residues. The remaining backbone models following these filtering steps contain long loops arranged in sheet-like structures ready for installation of additional sidechain based buttressing interactions.

We designed sequences onto these backbones focusing on further loop stabilization through buttressing (Fig. 1 bottom row). We began by scanning each position on the long loops for Asn, Asp, His or Gln placements that form backbone-sidechain bidentate hydrogen bonds between loops or between a loop and a helix, and for Val, Leu, Ile, Met and Phe placements that form loop-helix hydrophobic contacts; amino acids meeting these criteria were kept fixed in subsequent design steps. We then performed four rounds of full combinatorial Rosetta protein sequence design with slowly ramped up fa_rep weight to promote core packing; a slight compositional bias towards proline was used in the long loop to increase rigidity. The design models were filtered in Rosetta by number of buried unsatisfied heavy atoms (<=3), core residue hole score (<=−0.015), total score per residue (<=−2), packstat (>=0.5), and average hydrogen bond energy per residue (<=−1) in the buttressed long loops. The rigidity of the design models was evaluated using molecular dynamics simulations, and the extent to which the designed sequence encodes the structure by AlphaFold^12,13^. The *in-silico* validated designs span a diverse range of shapes with different repeat protein curvatures and loop geometries (Fig. 1B), and multiple loop buttressing strategies utilizing loop-helix hydrogen bond networks and loop-loop bidentate hydrogen bonds (Fig. 1C). These designed buttressed loops have significantly more diverse structures than the long, hairpin loops in the native ankyrins (Fig. S1), and contain more backbone hydrogen bonds (Fig. S2).

We expressed 102 selected designs (which we call Repeat proteins with Buttressed Loops (RBLs)) in *E. coli* and purified them by His tag-immobilized metal affinity chromatography. 77 of the purified proteins were soluble (representative models shown in Fig. 2A), 52 were monodisperse and 46 monomeric, as indicated by multi-angle light scattering coupled with size exclusion chromatography (SEC-MALS) (Fig. 2B). 44 of these proteins showed expected alpha-helical circular dichroism (CD) spectrum at 25°C, remained at least partially folded at 95°C and recovered nearly all the CD signal when cooled down to 25°C (Fig. 2C). 14 designs were further validated by small-angle X-ray scattering (SAXS) (Fig. 2D, Fig. S3); the experimental scattering curves were in agreement with profiles computed from the design models.

**Fig. 2.**
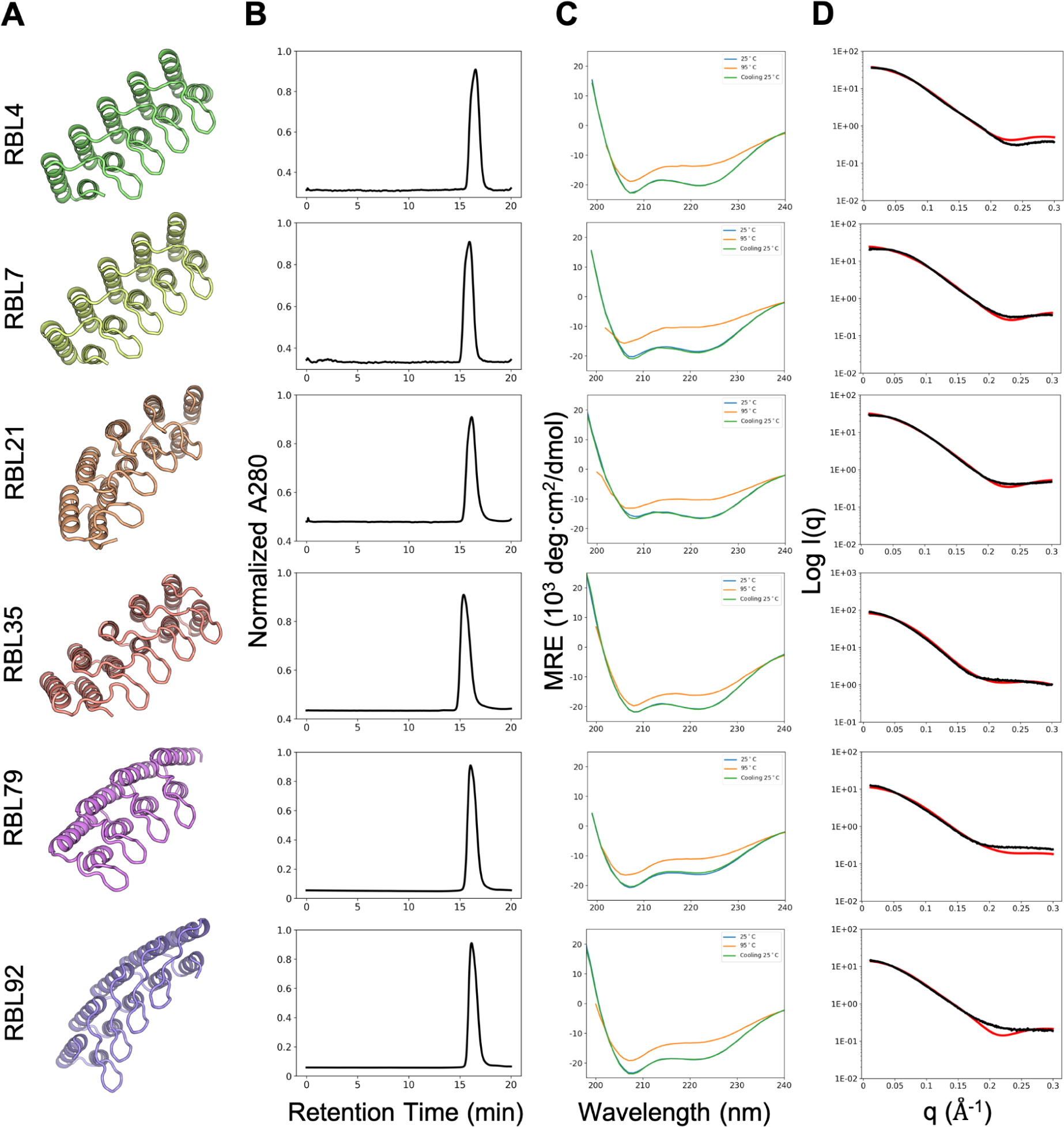
Biophysical characterization of designed helical repeat proteins with buttressed loops. **(A)** Structural models of six representative designs. **(B)** Traces of UV280 in size-exclusion chromatography. **(C)** Circular dichroism spectra collected at 25 °C (blue), 95 °C (orange) and 25 °C after heating (green). **(D)** Overlay of experimental (black) and theoretical (red) small angle X-ray scattering profiles.

We determined the crystal structure of design RBL4 at 1.8Å resolution (Fig3. A-E). RBL4 contains four helix-long-loop-helix repeat units that are sandwiched by two capping terminal capping helices. Each long loop is anchored on top of the neighboring helices and stabilized by interloop Asn-mediated bidentate hydrogen bonds networks as designed, and the design model is in good agreement with the crystal structure with a Cα RMSD of 1.7Å (Fig. 3A). The primary discrepancy between the crystal structure and design model is in the inter-repeat transformation: the design model is slightly curved (smaller superhelical radius) while the crystal structure is nearly flat (larger superhelical radius). Within individual repeat units, there is very close agreement between the crystal and design model, with repeat unit Cα RMSDs for different repeat units ranging from 0.48-0.61Å (Fig. 3B). The designed loop buttressing interactions–-the bidentate interloop hydrogen bonds (Fig. 3C) and loop-helix salt bridges (Fig. 3D)--were accurately recapitulated in the crystal structure. With the exception of the residues at the tips of the buttressed loops, the loop residues have B-factor values less than 60, suggesting they are reasonably well ordered (Fig. S4A).

**Fig. 3.**
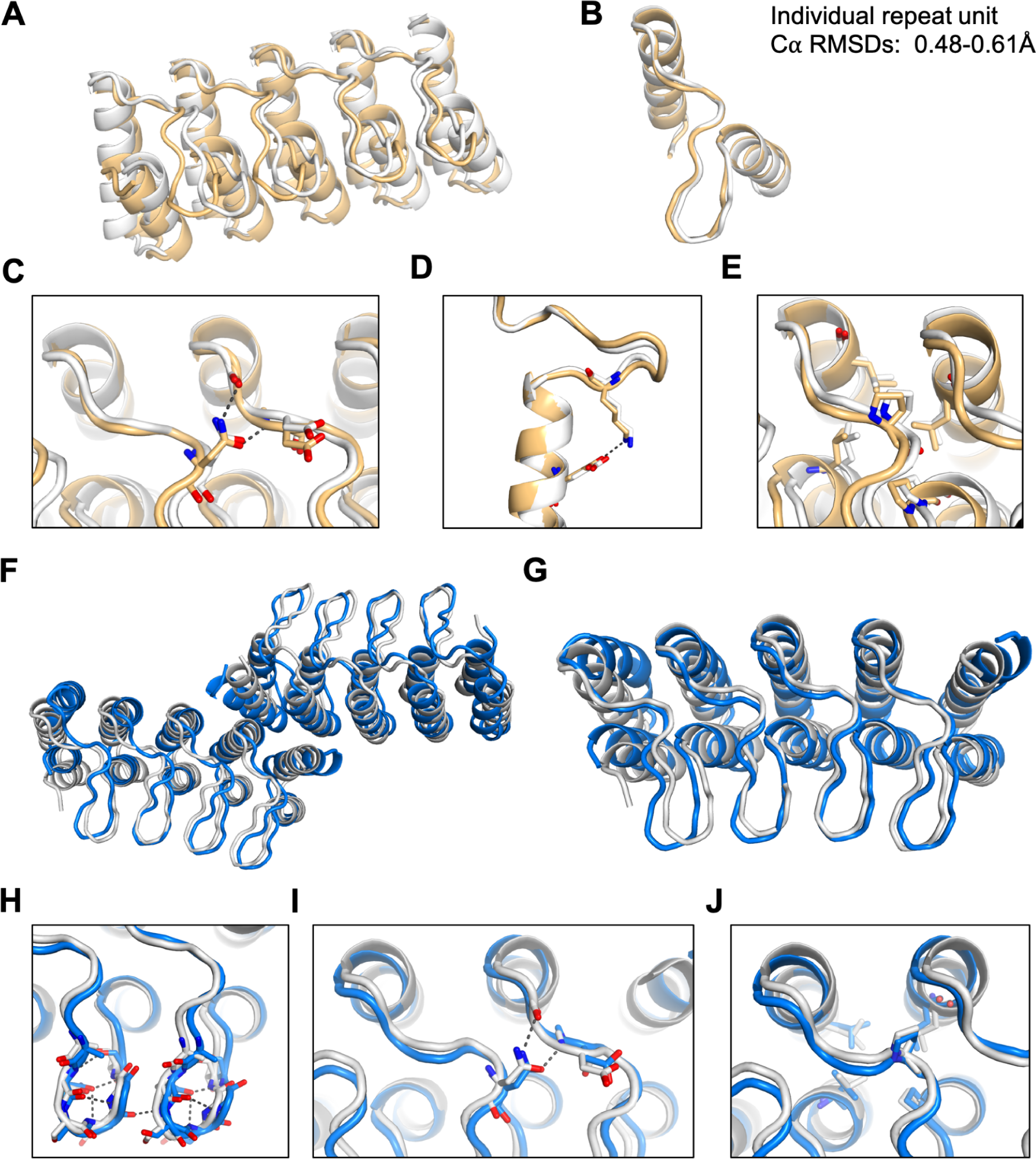
Structural characterization by X-ray crystallography. **(A)** Superimposition of crystal structure (yellow) onto the design model of RBL4 (gray). **(B)** Alignment of individual repeat units. **(C-E)** Accurately designed loop buttressing interactions. **(F)** Superimposition of crystal structure (blue) onto the design model of RBL7_C2_3 (gray). **(G)** Overlay of a monomer unit in the crystal structure onto the design model. **(H-J)** Accurately designed loop buttressing interactions.

Design RBL7 has similar overall geometry as RBL4 but with a smaller superhelical radius. This design was highly stable and monomeric, with an overall fold validated by SAXS (Fig. 2, second row). We obtained crystals which diffracted poorly with the highest resolution at 4.2Å. As previous studies suggested that synthetic oligomerization can sometimes assist crystallization^14^, we sought to generate a dimeric form of RBL7 by introducing a hydrophobic dimer interface. The redesigned protein, RBL7_C2_3, was soluble and dimeric, and we were able to solve the crystal structure at 3Å. The crystal structure closely matches the design model, with a Cα RMSD over the dimer of 2.9Å (Fig. 3F), and over the monomer, of 1.6Å (Fig. 3G). The main discrepancies between the crystal and designed structures were in the terminal helices. Similar to design RBL4, the crystal structure confirmed the accuracy of designed loop buttressing interactions in RBL7 (Fig. 3H-J). All of the designed interloop hydrogen bonds at the tip beta turns of long loops were recapitulated in the crystal structure (Fig. 3H). These hydrogen bonds likely are crucial for positioning the long loops and contribute to the close matching between the loops in the design model and those in the crystal structure. The B-factor values of the buttressed loops are slightly higher in RBL7_C2_3 (Fig. S4B) than in RBL4 (Fig. S4A), possibly because there are fewer loop stabilizing salt bridges analogous to those in RBL4 (Fig. 3D).

An exciting application of our designed RBLs is to use them as starting points for the computational design of high-affinity binding proteins. This could enable design of DARPin-like binders to a wide range of targets without the need for large scale library selection methods; the ability to design a wide diversity of repeating scaffolds with buttressed loops could considerably expand the space of targets. As a first step towards investigating the design of RBL based binders, we redesigned the extended groove bordered by the buttressed loops to bind extended peptides. To take advantage of the repeating nature of RBLs, we chose to focus on peptides with a repeating sequence motif–in this case, once a repeat unit is designed to bind a particular short peptide, repeat proteins containing multiple copies of this unit should bind peptides with multiple copies of the motif, provided the register between the repeat protein and the peptide can be maintained. Generalizing from the observation that some ankyrin family proteins can bind peptides with a PxLPxI/L (x can be any amino acid) sequence motif^15^, we sought to design binders for peptide sequences of the form XYZ_n_, where n is the number of repeats, X is a long polar residue interacting with residues in the buttressed loop beta turns, Y and Z are hydrophobic residues interacting with the helices and the helix-loop joint of RBLs (see Fig.4B for an example of one peptide repeat unit interacting with an RBL based peptide binder).

**Fig. 4.**
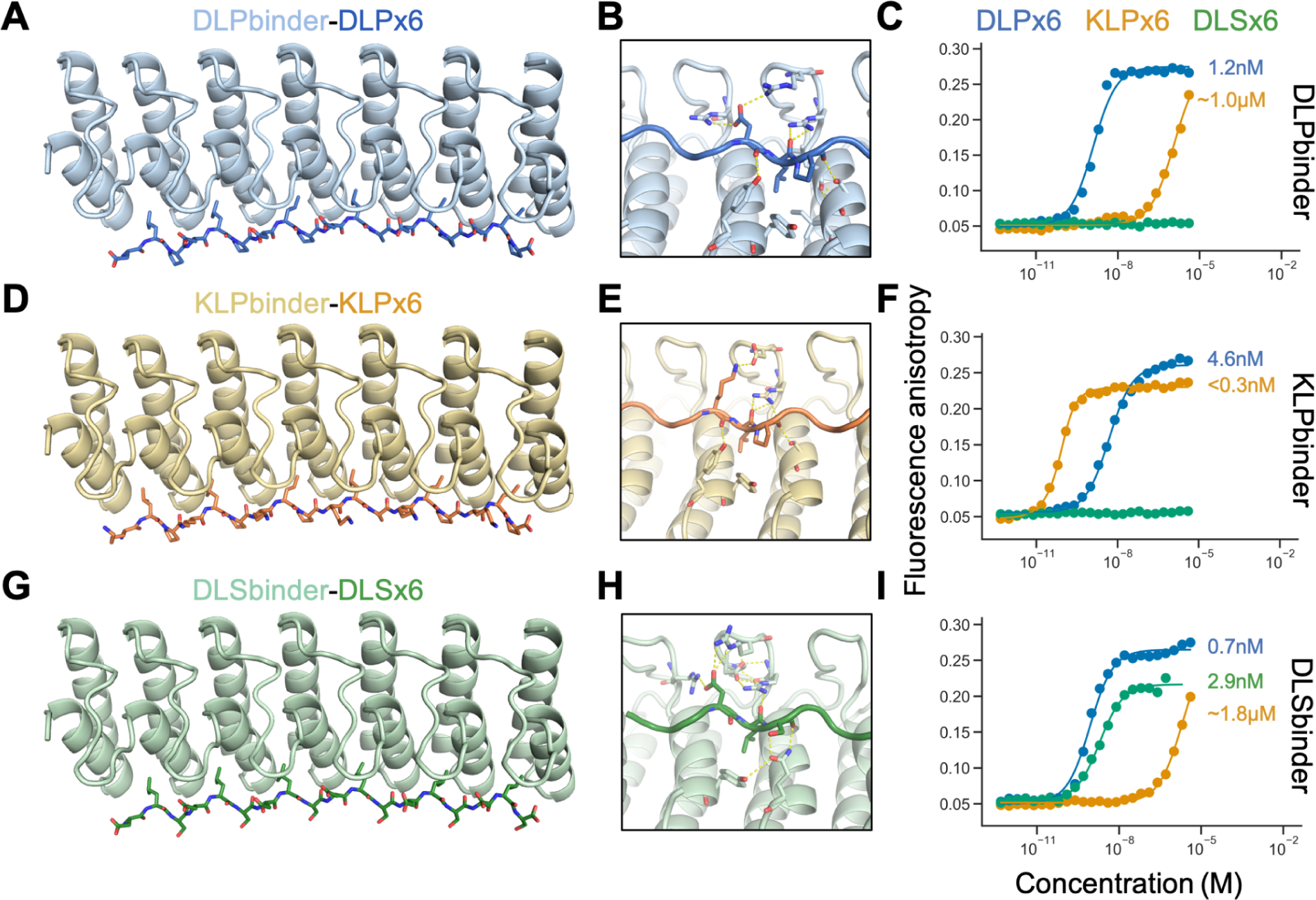
Designed repeat peptide binding RBLs. **(A, D and G)** Design models of binding proteins for peptides with 6 repeats of DLP, KLP and DLS respectively. **(B, E and H)** Sequence-specific interactions between the designed binders and the repeat peptides. **(C, F and I)** Fluorescence polarization measurement of binding of designs to repeat peptides. For each binder, a titration curve is plotted for the binding of each peptide (Blue: DLPx6, Orange: KLPx6, Green: DLSx6).

To design binders of XYZ_n_ peptides, we first docked tripeptide repeats in the polyproline II helix confirmation to the binding grooves of RBLs guided by the interactions in peptide-binding ankyrin family proteins in the PDB (see Methods), and carried out rigid body perturbations to diversify the docked poses. For each such pose, we used Rosetta sequence design to generate sequences of both RBL and peptide for optimized binding. We designed 34 proteins to bind six-repeat peptides (DLPx_6_, KLPx_6_ or DLSx_6_), screened them *in silico* based on protein-protein interaction design filters including AlphaFold^12,13^ structure recapitulation, obtained synthetic genes encoding the designs, and purified the proteins from *E. coli* expression. In initial binding screens using split luciferase assay, seven designs showed clear binding signals. From these designs, we selected the strongest binders for each peptide target, and performed fluorescence polarization measurements which showed the protein-peptide interactions are orthogonal and have high affinities (Fig. 4). All the selected binders were based on RBL4 with the peptides in nearly identical binding modes, with the Asp/Lys at the X position forming salt bridges with charged residues from the beta turn tip of RBL, the Leu at the Y position fully buried in the hydrophobic interface, and the Pro/Ser interacting with the residues on the bottom of helices (Fig. 4B, E and H). Both the DLPx_6_ binder and the KLPx_6_ binder bound their target peptides with high affinities (Kd=1.2nM and <0.3nM respectively) and high specificity (Fig. 4C and F). Neither binder bound DLSx_6_, suggesting Pro in the peptide was crucial in the protein-peptide interactions. We sought to rescue the DLSx_6_ binding by installing Gln-mediated bidentate hydrogen bonds (Fig. 4H). The resulting design bound DLSx_6_ with high affinity (Kd=2.9nM), but retained affinity for DLPx_6_ (Fig. 4I).

## Discussion

There are two primary routes forward for engineering new functions using our designed RBLs. First, by analogy with the many DARPins obtained starting from stabilized consensus ankyrin repeat proteins, it should be readily possible to create binders by random library generation in conjunction with yeast display and other selection methods for binding to targets of interest. Second, as suggested by our design of peptide binding proteins, computational design methods can be used to generate binders to a wide variety of targets taking advantage of the diverse geometries that can be achieved with different buttressed loops on different repeat protein scaffolds.

From a fundamental design perspective, the crystal structures presented here show that computational protein design has advanced to the point that proteins with multiple ordered long loops can now be designed. Key to this success were the design of dense networks of hydrogen bonding and nonpolar interactions within and between the loops, and between the loops and the underlying secondary structural elements. Our approach alone or integrated with additional recent progress in loop design^16^ and recently developed deep learning approaches for protein design^17–21^ (which do not currently address the challenge of designing structured long loops), should enable the design of structured loops for binding functions and beyond in a wide variety of scaffolds. For example, for enzyme design, multiple loops emanating from designed TIM barrels^22–24^ could be built to buttress each other and, together with the residues emerging from the top of the beta strands and helices in the TIM structure, form an extensive catalytic site and associated substrate/transition state binding site.

## Methods

### Computational design method

We developed our computational protein design protocols using Rosetta^25,26^ (2019.01) and PyRosetta4 (release 2019.22)^27^. Our protocol of parametric repeat protein generation started by building an ideal helix H1 (with the length of 12 to 28 residues) with MakeBundleHelix mover in Rosetta^25,26^, and placing it away from the Z axis with a given radius and an angle corresponding to its orientation. A second helix, H2 (with the length of 12 to 28 residues), was then modeled and placed according to the specification of the six rigid-body degrees of freedom for geometry transformation from H1 to H2. By combining H1 and H2 into one pose, we built the first repeat unit R1. Subsequently, we used user-specified six rigid-body degrees of freedom between repeat units to perform geometric transformation to obtain the second unit R2. Based on the number of repeats desired, we propagated the repeat units to generate the helical repeat protein backbones. We then connected pairs of sequence-adjacent helices with loops of 3-6aa using ConnectChainMover^10^. To filter generated repeat protein backbones, we required that the termini of the helices to be connected by buttressed long loops have to be less than 18Å apart. We also removed the low-quality backbone models with fewer than 28% of the residues in a buried core.

To design buttressed loops, we developed a hybrid method that assembles native structural motifs via kinematic loop closure. To guide the sampling towards the hairpin-shape conformations, we constructed a motif library that consists of native beta turns. A beta turn motif is defined by having a backbone-to-backbone hydrogen bond between the carbonyl group of residue i and the amine group of residue i+3^28,29^. In this work, we searched for native beta turn fragments by mining a set of selected PDB based on 90% maximum sequence identity and 1.6 Å resolution cutoff from PISCES^30^. The collected beta turns were further clustered by K-centers algorithm^31^ at maximum cluster distance of 0.63Å, resulting in 180 motif clusters. Using the same approach, we compiled a library of native helical capping motifs to guide the sampling of loops connecting helices in the repeat proteins.

We used GeneralizedKIC (GenKIC)^11^ for loop closure. An extended loop fragment was first constructed by stitching native helical capping motifs (4aa), beta turn motifs (4aa) and KIC residues (5-10aa) with randomized backbone torsion angles. We chose these lengths because we found limited structural diversity for loops with length less than 9aa. When the loop length exceeded 14aa, it became significantly difficult to design buttressing interactions to stabilize the entire loop. The torsion angles of beta turns were set according to the motifs sampled from the beta-turn library; and the phi/psi torsion angles of non-pivot KIC residues were sampled from Ramachandran distribution, with omega torsion angles fixed at 180 degrees. All the bond lengths were kept fixed at the ideal lengths. The position of the beta turn was randomly sampled in the loop. In each step of GenKIC, kinematic loop closure was performed to connect the loop to the intended insertion site. Loop conformations were filtered by backbone steric clashes. We further filtered the models by selecting loops with at least two intraloop backbone-to-backbone hydrogen bonds. To avoid the helical conformations, we removed the models that are predicted to have more than 5 consecutive helical residues by DSSP^32^. This ensured the extended beta-hairpin shape which contributed to the loop stability and compatibility for buttressing.

To install the loops of the same conformation in each unit of repeat proteins, we used the RepeatPropagationMover in Rosetta^25,26^. After filtering out the loops with steric clashes, we computed three metrics to help select the best loop conformations for buttressing: number of interloop backbone-to-backbone hydrogen bonds, loop motif scores and direction scores. We required at least one interloop backbone-to-backbone hydrogen bond between each pair of neighboring loops to maximize the sequence-independent loop buttressing effect. To select loops with loop-helix hydrophobic contacts, the motif scores were computed by matching the selected pairs of residues to the known native hydrophobic residue pairs (Val, Leu, Ile, Met and Phe) in PDB^33^. The scores for each match of residue pairs in the loop regions were then summed to one total score. Only the loops with a negative total motif score were selected. The direction scores described the relative orientation of the loops from the rest of input repeat proteins. Specifically, we defined two vectors: vector **a** started from the center of mass of the two loop terminal residues, to the farthest Calpha atom of the loop; vector **b** started from the same point as **a**, but pointed towards the center of mass of the repeat unit. The direction score was derived by computing the angle between the two vectors:

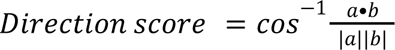

The accepted angles ranged from 45° to 135°. We also required at least five residues within 8Å of the closest helical residues.

Next, we performed a fast sequence design task to identify loop conformations compatible with interloop bidentate hydrogen bond networks. From each propagated set of loops, the loop on the second repeat unit was selected for sequence design. One packing step using PackRotamersMover^25,26^ was conducted separately for each residue on this loop using amino acids that are compatible with forming side-chain-to-backbone bidentate hydrogen bonds: Asn, Asp, Gln and His. We excluded amino acids with long side chains (Arg and Lys), as their high entropic cost might diminish the free energy contribution of buttressing. After each packing step, bidentate hydrogen bonds between the packed residue and its neighboring residues were counted. Specifically, a bidentate hydrogen bond was defined as two separate hydrogen bonds forming between atoms in the functional group of the side chain and the atoms in the backbone of the neighboring repeat unit. The selected amino acid was kept only if it formed interloop bidentate hydrogen bonds; otherwise, the original amino acid (by default Ala) was kept. In the case where the one-step packing approach failed to generate any interloop bidentate hydrogen bonds, we used an alternative three-stage scheme to maximize the sampling efficiency of bidentate hydrogen bonds: identifying pseudo bidentate hydrogen bonds, constrained minimization for building hydrogen bonds and evaluating the resulting bidentate hydrogen bonds. We defined that a pseudo hydrogen bond has a donor-acceptor distance < 3Å and a hydrogen bond angle > 120°. After propagating the designed residue to all the repeat units, we imposed a harmonic distance constraint between each donor and acceptor atoms with target distance as 2Å and standard deviation as 0.5Å. At the minimization stage, we performed symmetric minimization of the loops to improve the interactions of potential hydrogen bonds. Finally, using the Rosetta score function, we searched for bidentate hydrogen bonds in the minimized loop conformations.

To guide the sequence design, we used LayerSelector to define the core, the boundary and the surface layers, and specified the allowed amino acids for each layer. We added residue type constraints to fix the identity of the residues participating loop buttressing bidentate hydrogen bonds, so the stabilizing interactions obtained during loop sampling would be maintained throughout sequence design. Next, we performed four rounds of sequence design using FastDesign mover under the repeat symmetric constraints to ensure the repeat units had the same structures and sequences. To improve the solubility and folding of the designs, we subsequently performed one round of FastDesign for the solvent-exposed hydrophobic residues on the terminal repeat units. Only polar residues such as Glu, Gln, Lys and Arg were allowed for this round of design.

The designed structures were then refined by minimization in Cartesian space and subsequently filtered by number of buried unsatisfied heavy atoms (<=3), hole score normalized by total number of core residues (<=−0.015), total score normalized by total number of residues (<−2), packstat (>=0.5) and hydrogen bonding energy of each loop residue (<=−1). Top 10% scoring structures were further tested by *in silico* validation methods such as molecular dynamics simulations (Ca RMSD < 3Å), or AlphaFold^12,13^ (PLDDT > 80, RMSD < 3Å) or RoseTTAFold^34^ (PLDDT > 80, RMSD < 3Å). Structural similarity between native ankyrin loops and the designed RBL loops were computed by TM-align^35^.

We performed molecular dynamics simulations using GROMACS 2018.4^36^ with the Amber99SB-ILDN force field^37^. The design models were solvated in dodecahedron boxes of the explicit TIP3P^38^ waters with the net charge neutralized. We treated long-range electrostatic interactions with the Particle-Mesh Ewald method^39^. Both short-range electrostatic interactions and van der Waals interactions used a cutoff of 10Å. Energy minimization was performed using the steepest descent algorithm. A 1-ns equilibration under the NPT ensemble was subsequently performed with position restraints on the heavy atoms. We used Parrinello–Rahman barostat^40^ and Velocity-rescaling thermostat^41^ for pressure coupling (1 atm) and temperature coupling (310K) respectively. For the production runs, we launched three 20-ns trajectories under NPT ensemble for each design model. The Ca atom RMSD against the design model was computed for analysis.

### Protein expression and characterization

Genes encoding the *in silico* validated designs were synthesized (IDT) and cloned into pET-29b expression vectors. The plasmids were transformed into Lemo21 (DE3) expression *E. coli* strain (NEB). Protein expression was performed using auto-induction protocol^42^ at 37°C for 24h in 50ml or 100ml culture. During the purification, cells were pelleted at 4,000g for 10min and resuspended in 25ml lysis buffer (25mM Tris-HCl pH=8, 150mM NaCl, 30mM imidazole, 1mM DNase and 10mM lysozyme with Pierce Protease Inhibitor Tablets (Thermo Fisher)). Sonication was subsequently performed for 2.5 min (10s on and 10s off per cycle). The lysate was then centrifuged at 16,000g for 30 min. The supernatant was applied to a gravity flow column packed with Ni-NTA resin (Qiagen), followed by 20ml wash buffer (25mM Tris-HCl pH=8, 150mM NaCl, 30mM imidazole) and 5ml elution buffer (25mM Tris-HCl pH=8, 150mM NaCl, 400mM imidazole). The eluted protein was then concentrated and injected into an Akta Pure FPLC device with the flow rate of 0.75ml/min in the running buffer (25mM Tris-HCl pH=8, 150mM NaCl). The typical yield of a monodisperse and thermally stable designed RBL is 1-6g/L. To perform multi-angle light scattering coupled with size exclusion chromatography, we prepared the purified protein at ∼2mg/ml and injected 100μl of sample into a Superdex 200 10/300GL column and measured the light scattering signals using a miniDAWN TREOS device (Wyatt Technology). To measure the circular dichroism (CD) signals, we first prepared the sample at ∼0.2mg/ml in 25mM phosphate buffer in 1mm cuvette. A Jasco J-1500 CD spectrometer was used for all CD measurements. We set the range of wavelength from 190nm to 260nm and scanned over a three-temperature (25°C, 95°C and cooling back at 25°C) set for each sample. We submitted all samples for small-angle X-ray scattering (SAXS)^43,44^ to Advanced Light Source, LBNL for data collection at the SIBYLS 12.3.1 beamline.

### Design and characterization of repeat peptide binding proteins

We used the recently developed protein interface design method^7^ for *in silico* binder docking and design experiments. Docking of repeat peptides to the binder scaffold was guided by the geometric transformation between native ankyrins to their peptide targets in the crystal structures from PDB^15^. Symmetric sequence design was performed for each docked peptide-protein pair following the same protocol used for designing RBLs. All the designed complexes were computationally tested by AlphaFold with a cutoff of PAE_interaction<=15 before experimental characterization.

Split-luciferase assay was performed using the Nano-Glo Luciferase Assay System (Promega). The coding sequence of small-BiT was fused to the gene of peptide binders and the coding sequence of large-BiT was fused to the coding sequence of the target peptide (Genscript). The BiT-fused proteins and peptides were expressed and purified with the same protocol for RBLs. The purified peptide binders and target peptides were titrated in the presence of Nano-Glo substrate in 96-well plates and the luminescence was measured on a Synergy Neo2 plate reader (Agilent). To conduct the fluorescence polarization binding assays, we synthesized the repeat peptide fragments with 5’-TAMRA labels. Fluorescence polarization measurements were performed at 25°C in a Synergy Neo2 plate reader (Agilent) with a 530/590nm filter. Series of two-fold dilutions of binder-peptide 80-ul mixture were performed in 25mM Tris-HCl pH=8, 150mM NaCl, 0.05% v/v TWEEN20 in 96-well assay plates. The protein concentrations ranged from 4uM to 0.47pM and the concentration of TAMRA-labeled peptide was kept at 0.3nM. The samples were incubated for 3h before measurement.

### Structural characterization by X-ray crystallography

RBL4 was concentrated to 150 mg/ml and crystallized by vapor diffusion. Initial crystals formed in the MCSG-2 crystallization screen (Anatrace) and optimized crystals were grown in 100 mM sodium acetate, pH 4.4, and 2% PEG 4000. The crystal was cryoprotected with 30% ethylene glycol and flash cooled in liquid nitrogen. Diffraction was measured at APS beamline 23 ID-B. Reflections were indexed, integrated, and scaled with autoproc^45^. The structure was solved by molecular replacement in Phaser^46^. Initial attempts using the predicted model were unsuccessful due to clashes. A subsequent search for 8 copies of a single helix-loop-helix repeat (residues 76-118) identified two copies of the protein in the asymmetric unit. The model was rebuilt using phenix autobuild^47^ and completed by iterative rounds of interactive refinement in coot^48^ and reciprocal space refinement in phenix^49–52^. The final refinement strategy included reciprocal space refinement, individual ADPs, TLS refinement using parameters determined with TLSMD^53^, and occupancy refinement of alternate conformations. Model geometry was assessed with Molprobity^54^. The final model included 99.5% of residues in the favored region of the Ramachandran plot with no outliers.

RBL7_C2_3 was concentrated to 119 mg/ml and crystallized by vapor diffusion in 2.4 M sodium malonate, pH 7.0, using the MCSG-1 crystallization screen (Anatrace). The crystal was cryoprotected by addition of 10 volumes of 3.4 M sodium malonate, pH 7.0 and flash cooled in liquid nitrogen. Reflections were indexed, integrated, and scaled with XDS^55^. To solve the structure by molecular replacement, an ensemble of monomer structures was generated by AlphaFold and used as a search ensemble in Phaser. The solution contained 8 molecules that formed 4 homodimers. The model was rebuilt with phenix autobuild with morphing and completed by iterative rounds of interactive refinement in coot and reciprocal space refinement in Buster^56^ or phenix. The final refinement strategy in phenix included reciprocal space refinement, individual ADPs, NCS restraints, and TLS refinement using one group per chain. The final model had 98.22% of residues in the favored regions of the Ramachandran plot with no outliers. Crystallographic software was installed and maintained using SBGrid^57^.

**Table 1.** Crystallographic data collection and refinement statistics.

|  | **RBL4** | **RBL7_C2_3** |
| --- | --- | --- |
| **Wavelength (Å)** | 1.033 | 0.9795 |
| **Resolution range (Å)** | 59.84 - 1.8 (1.864 - 1.8) | 45.3 - 2.986 (3.092 - 2.986) |
| **Space group** | P 21 21 2 | P 21 21 21 |
| **Unit cell (Å, °)** | 62.287 215.442 28.008 90<br>90 90 | 111.961 117.644 142.056 90<br>90 90 |
| **Total reflections** | 258923 (25447) | 258197 (25212) |
| **Unique reflections** | 35977 (3521) | 38693 (3751) |
| **Multiplicity** | 7.2 (7.2) | 6.7 (6.7) |
| **Completeness (%)** | 98.67 (96.09) | 99.02 (92.46) |
| **Mean I/sigma(I)** | 5.70 (0.36) | 5.34 (0.64) |
| **Wilson B-factor** | 40.77 | 67.39 |
| **R-sym** | 0.1381 (3.305) | 0.3314 (3.08) |
| **R-meas** | 0.1498 (3.557) | 0.3597 (3.336) |
| **R-pim** | 0.05685 (1.298) | 0.1385 (1.269) |
| **CC1/2** | 0.989 (0.335) | 0.996 (0.395) |
| **Reflections used in refinement** | 35803 (3439) | 38424 (3544) |
| **Reflections used for R-free** | 1974 (193) | 1988 (179) |
| **R-work** | 0.2329 (0.6169) | 0.2097 (0.3431) |
| **R-free** | 0.2835 (0.6811) | 0.2593 (0.3912) |
| **Number of non-hydrogen atoms** | 3216 | 12376 |
| **macromolecules** | 3066 | 12331 |
| **ligands** | 28 | 7 |
| **solvent** | 122 | 38 |
| **Protein residues** | 410 | 1648 |
| **RMS(bonds) (Å)** | 0.011 | 0.004 |
| **RMS(angles) (°)** | 1.07 | 0.70 |
| **Ramachandran favored (%)** | 99.51 | 98.35 |
| **Ramachandran allowed (%)** | 0.49 | 1.65 |
| **Ramachandran outliers (%)** | 0.00 | 0.00 |
| **Rotamer outliers (%)** | 0.96 | 3.80 |
| **Clashscore** | 2.22 | 7.38 |
| **Average B-factor (Å<sup>2</sup>)** | 52.05 | 77.82 |
| **macromolecules** | 52.07 | 77.86 |
| **ligands** | 51.18 | 121.17 |
| **solvent** | 51.77 | 55.70 |
| **Number of TLS groups** | 4 | 8 |

Statistics for the highest-resolution shell are shown in parentheses.

## Data availability

All the design models, protein sequences and DNA sequences are available at: https://files.ipd.uw.edu/pub/2023_buttressed_loops/data.tar.gz. Crystal structures and reflection data have been deposited in the RCSB Protein Data Bank with accession ids 8FRE (RBL4) and 8FRF (RBL7_C2_3). X-ray diffraction images have been deposited in the SBGrid Data Bank.

## Code availability

The design scripts for parametric repeat protein generation and buttressed loop design are available at: https://github.com/hanlunj/buttressed_loops.git.

## Acknowledgements

We thank F. Praetorius, P. Leung and S. Vazquez for advice on fluorescence polarization assay; B. Wicky and I. Lutz for advice on split luciferase assay; the Wysocki group at Ohio State University for the support with native mass spectrometry; ALS SIBYLS beamline at Lawrence Berkeley National Laboratory for SAXS data collection; K. VanWormer for wet lab support; L. Goldschmidt for computing support; D. Silva, C. Xu, H. Bai, C. Norn, P. Salveson, D. Sahtoe, R. Kibler, B. Weitzner, F. DiMaio, P. Bradley, B. Stoddard, K. Lee and F. Pardo for helpful discussions. Funding for this work is provided by the Audacious Project at the Institute for Protein Design and the Open Philanthropy Project Improving Protein Design Fund. Use of the Stanford Synchrotron Radiation Lightsource, SLAC National Accelerator Laboratory, is supported by the U.S. Department of Energy, Office of Science, Office of Basic Energy Sciences under Contract No. DE-AC02-76SF00515. The SSRL Structural Molecular Biology Program is supported by the DOE Office of Biological and Environmental Research, and by the National Institutes of Health, National Institute of General Medical Sciences (P30GM133894). GM/CA@APS has been funded by the National Cancer Institute (ACB-12002) and the National Institute of General Medical Sciences (AGM-12006, P30GM138396). This research used resources of the Advanced Photon Source, a U.S. Department of Energy (DOE) Office of Science User Facility operated for the DOE Office of Science by Argonne National Laboratory under Contract No. DE-AC02-06CH11357. The Eiger 16M detector at GM/CA-XSD was funded by NIH grant S10 OD012289.

---

**Fig. S1.**
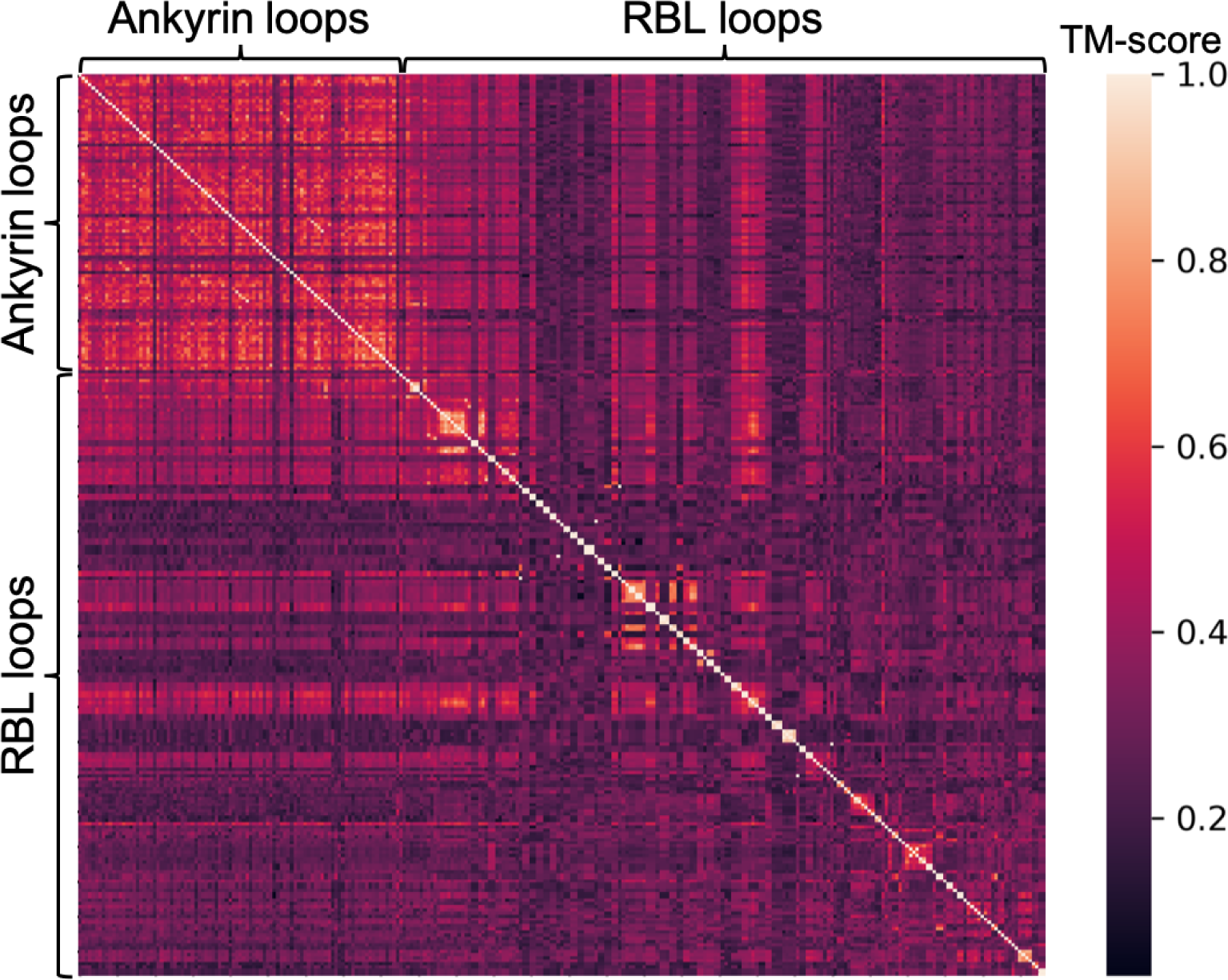
TM-scores between native ankyrin loops and the designed loops from RBLs. Higher TM-scores indicate higher structural similarity.

**Fig. S2.**
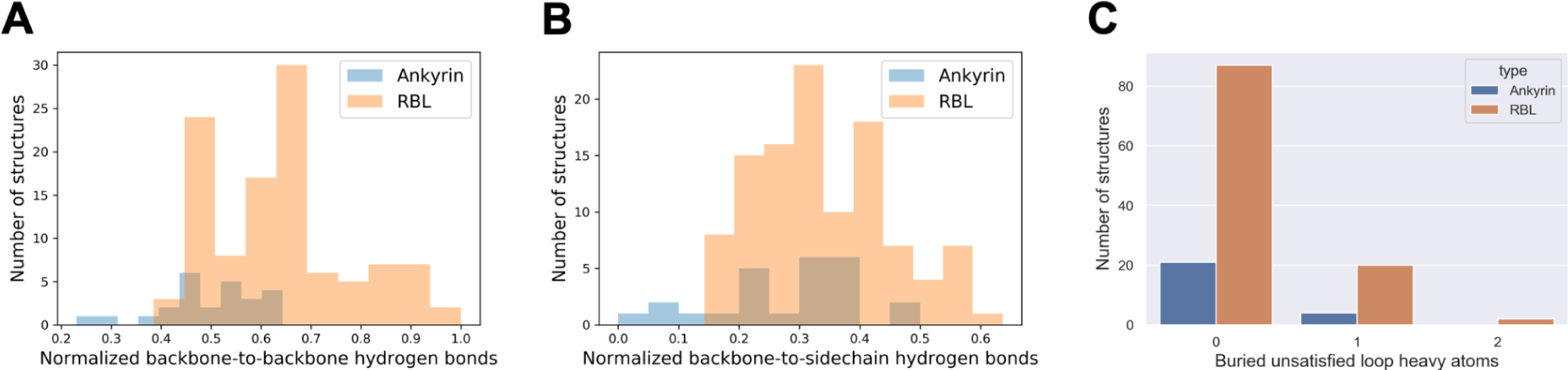
Comparison of hydrogen bonds and buried unsatisfied loop heavy atoms between ankyrin loops and the designed loops in RBLs. **(A)** Distribution of the number of backbone-to-backbone hydrogen bonds involving one long loop in each structure normalized by the loop length. **(B)** Distribution of the number of backbone-to-sidechain hydrogen bonds involving one long loop in each structure normalized by the loop length. **(C)** Number of buried unsatisfied loop heavy atoms in one long loop in each structure.

**Fig. S3.**
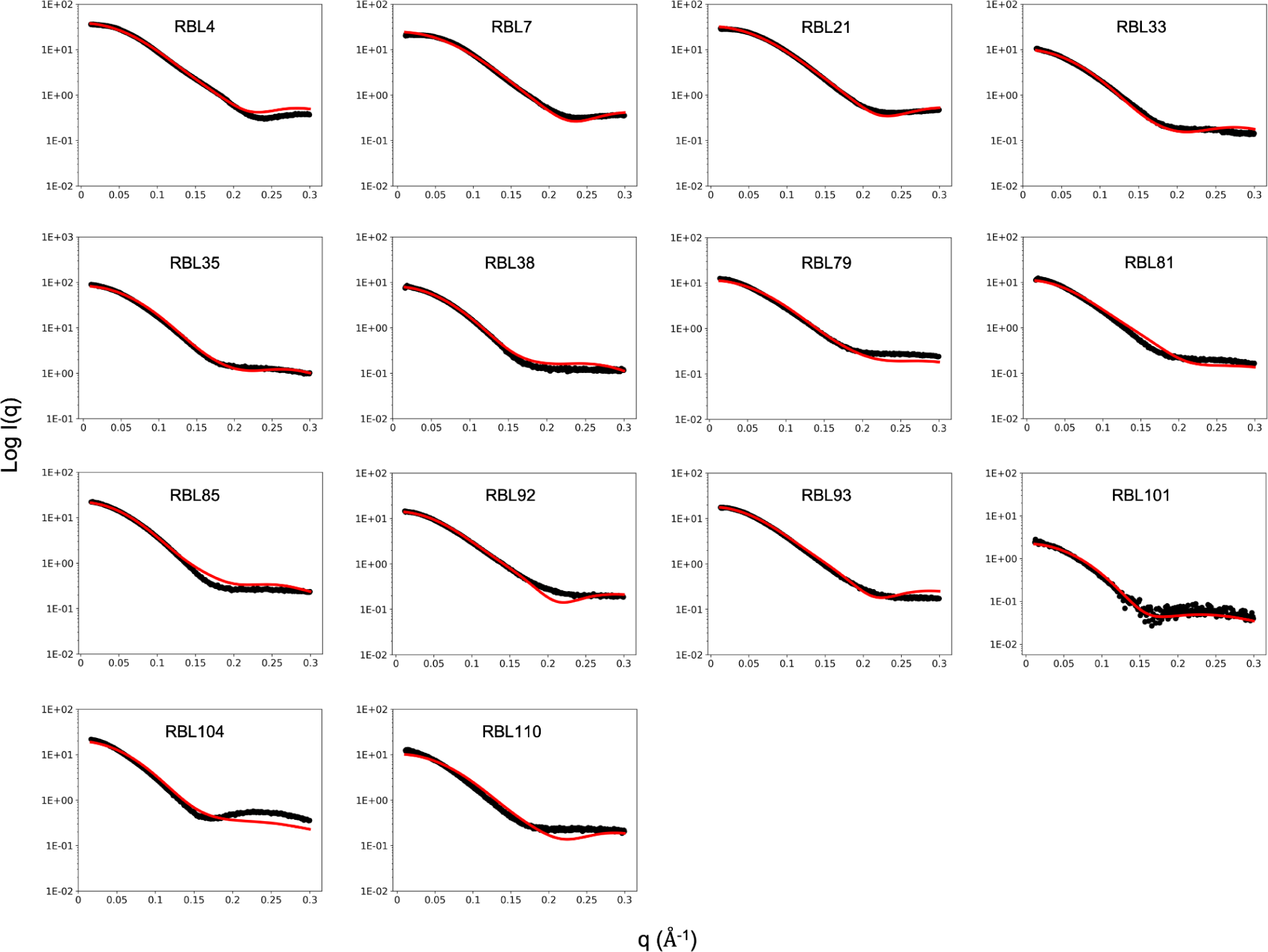
Overlay of experimental (black) and theoretical (red) small angle X-ray scattering profiles.

**Fig. S4.**
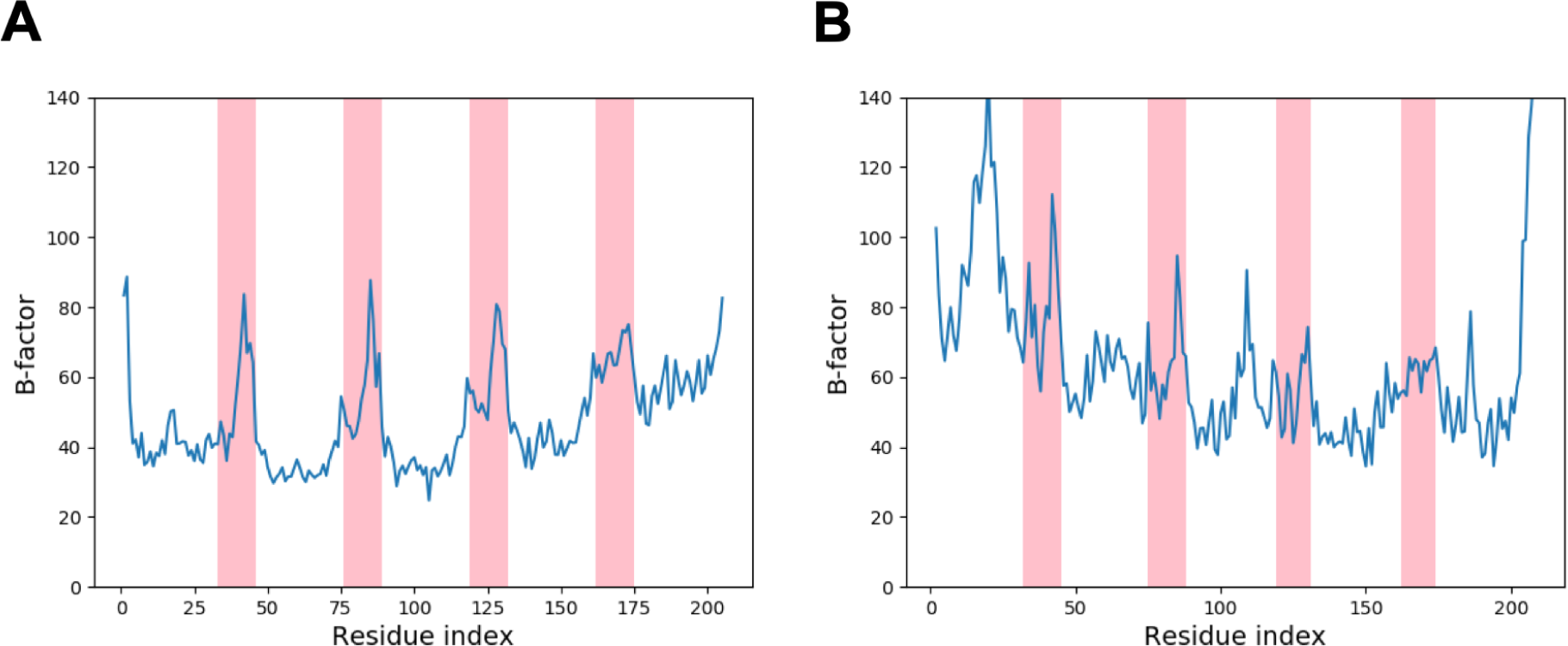
Residue-wise B-factor values of the crystal structures. **(A)** RBL4. **(B)** RBL7_C2_3. The regions corresponding to the buttressed long loops are highlighted in pink.

